# Precision Sub-phenotyping of Histologically Stable Kidney Transplants by a Composite Molecular and Cellular Instability Score

**DOI:** 10.1101/2019.12.18.881706

**Authors:** D. Rychkov, S. Sur, M. Sirota, M. M. Sarwal

## Abstract

Acute Rejection (AR) is the main cause of the graft dysfunction and premature graft loss, and diagnosis of rejection before advanced histological injury is crucial to salvage graft function. However, recent molecular studies have highlighted the unrecognized burden of sub-clinical graft rejection when graft function is preserved, and a dichotomy exists, of a histologically normal biopsy with molecular signatures of AR. Conversely, significant variation also exists in the definition of a stable allograft, defined as a transplant with absence of clinical AR, with absence of histological inflammation, though published studies have highlighted that some of these stable samples will not have stable immune quiescence, as they may be molecularly similar to AR. Thus, refining the definition of a stable allograft as one that is clinically, histologically *and* molecularly quiescent is critical, as the inclusion of stable allografts in mechanistic and clinical studies are vital to provide a normal, non-injured comparative group for all interrogative studies on understanding allograft injury.

With this goal in mind, we analyzed publicly available transcriptional data across 4,845 human kidney tissue samples from 38 Gene Expression Omnibus (GEO) datasets, inclusive of 510 allograft biopsy samples with AR, 1,154 renal allograft biopsies classified in each dataset as histological stable (hSTA), and 609 normal kidney (donor) samples. By applying a machine learning model, a substantive number of hSTA samples were found to be molecularly similar to AR (mAR); these have been reclassified in this study as clinical and histological stable samples with transcriptional signatures overlapping with AR (hSTA/mAR), with the predominant expression of a subset of 6 genes (*KLF4, CENPJ, KLF2, PPP1R15A, FOSB, TNFAIP3*). To understand the cellular sources of these molecular signals, we utilized xCell, a cell type enrichment tool and interrogated 64 specific cell types to identify 5 (*CD4+ Tcm, CD4+ Tem, CD8+ Tem, NK cells*, and *Th1 cells*) that were also highly predictive for classification of the AR phenotype in these studies. A combined gene and cell-type specific *InstaScore* (AUC 0.99) was developed using gene and cell subtype data to re-phenotype all hSTA allografts. This clearly defined two disparate hSTA biopsies: those that are both histologically and molecularly quiescent (hSTA/mSTA) or those that are histologically quiescent but molecularly similar to AR (hSTA/mAR). The clinical utility of the Instability Score was subsequently assessed by independent validation on a serial set of post-transplant hSTA biopsies, where strong significant correlation was observed between the score on 6 month post-transplant hSTA graft biopsies, where hSTA/mAR samples had a significant change in graft function and graft loss at 5 year follow-up.

In conclusion, our computational approach of precision sub-phenotyping of hSTA allografts by the *InstaScore* identifies discrepancies in the current recognition of a stable allograft by histology alone. Precision molecular sub-typing of the hSTA allograft into the hSTA/mSTA group is an important deliverable for selection of “true” STA samples for mechanistic studies, and into the hSTA/mAR group, for accurate prediction of subsequent patient clinical outcomes, and real time treatment stratification for hSTA/mAR allografts to positively impact long-term graft survival.

## Introduction

Breakthroughs in surgical approaches for organ transplantation and development of newer generations of immunosuppressive drugs to support engraftment across HLA barriers have resulted in a dramatic reduction in mortality among patients with an end stage organ disease, improvements in life quality and cognition, over the last decade[1]. However, overt and covert allograft acute rejection (AR) remains the most functionally relevant immune escape that drives adverse graft outcomes. Recent advances in omics technologies have uncovered the hidden burden of subclinical acute rejection (sAR), which can be observed even in the presence of an otherwise histologically pristine allograft [2,3]. Thus, precision molecular tissue phenotyping is needed to fully understand the granular phenotypes of clinical, histological and molecular injury from AR and to map clinical, histological and molecular quiescence for the true classification of a histologically stable (hSTA) allograft.

In this study, we propose that precision phenotyping of a kidney transplant biopsy by applying machine learning to gene expression measures which can enhance the accuracy of biopsy diagnosis [4]. The large amount of various molecular data in the public domain: genomic, transcriptomic, proteomic, and metabolomic, spurred by the expansion of next-generation sequencing technologies, coupled with advances in analytical and statistical methods, has already triggered the acceleration of the biological research in the field of personalized treatment and precision phenotyping in many other biomedical fields [5–7]. It now allows a unique opportunity to re-use aggregated human organ transplant related datasets in the Gene Expression Omnibus (GEO) [8], regardless of technology and platform era differences, utilizing novel approaches for data normalization and meta-analyses. Our group pioneered the meta-analysis of human biopsy transplant cohorts from four different solid organs - heart, kidney, liver, and lung – to identify and validate a common rejection module (CRM) of 11 genes (*BASP1*, *CD6*, *CXCL10*, *CXCL9*, *INPP5D*, *ISG20*, *LCK*, *NKG7*, *PSMB9*, *RUNX3*, and *TAP1*) that were significantly overexpressed in AR across all tissues, irrespective of organ source[9]. A similar approach utilized for 7 cross-organ tissue datasets confirmed a similar signature for AR (*ISG20*, *CXCL9*, *CXCL10*, *CCL19*, *FCER1G*, *PMSE1*, and *UBD)* [10], and 6 kidney transplant biopsy datasets were interrogated to identify an 85-gene signature associated with interstitial fibrosis and tubular atrophy [11]. Other meta-analysis studies on public data have been performed to study peripheral blood signatures of AR [12].

A significant obstacle in mining public transplant data is the difficulty in assigning what a truly stable transplant is, an issue of critical importance in this kidney of analysis, as this group of samples forms the control, “no-injury”, immune-quiescent group for the analysis. Not only is there significant discrepancy (19-55%) among pathologists for histological phenotyping [13,14], the histologic grading in renal transplant pathology is made within the Banff classification [15] which covers 10 different histologic lesions with an empirically derived, subjective scale of 0 to 3, and alloantibody assessment and monitoring varies widely between institutions [16], resulting in considerable disagreements in the histological interpretation of what is a stable graft [17]. In a recent study [18], unsupervised archetypal analysis and reclassification of 1,208 biopsy samples resulted in 32% disagreement of transcriptional phenotypes with histological diagnosis [13,14]. In addition, work from our group and others [2,3,18–22] has confirmed that sub-clinical molecular inflammation and AR related transcriptional and protein signatures can exist in absence of obvious infiltrates by histology, and can predate full blown histological rejection and increase the risk of progressive chronic graft injury. Thus, the ability to truly define a “stable” allograft becomes a critical unmet need for clinical assessment of the allograft and for the conduct of interrogative mechanistic studies into allograft pathology and injury. Simply accepting all samples classified as “stable” from different investigators and datasets will neglect the important biological diversity that naturally exists in the histological definition of a “stable” sample and will necessarily lead to false positive and false negative discoveries when attempting to study graft rejection.

We hypothesize that accurate classification of a stable allograft is not only a critical requirement for the conduct of successful meta-analyses of transplant injury, but also, more precise (sub-)phenotyping and early recognition of AR of otherwise functionally and histologically stable allografts on protocol biopsies, will allow timely and proactive adjustment of immunosuppression to immunologic risk, with a positive impact on graft survival and the patient’s quality of life [5,17]. To enable this, we have leveraged the largest public dataset analyzed in human renal transplantation: 4,845 kidney tissue microarray samples from 38 publicly available human normal kidney and transplant kidney tissue datasets, run on 18 different microarray platforms, to investigate the molecular diversity of histologically stable allografts. This study provides a new composite gene and cell specific analysis tool, the Instability Score (or *InstaScore*), that could provide an important adjunct to a pathologist for comprehensive and highly quantitative phenotyping of a protocol kidney transplant biopsy, in addition to the standard histological assessment.

## Methods and Materials

### Data collection

We carried out a comprehensive search for publicly available microarray data at NCBI GEO (Gene Expression Omnibus) database [8] (http://www.ncbi.nlm.nih.gov/geo/). We used the keywords “kidney”, “renal”, “transplant”, and “biopsy”, among organisms “Homo Sapiens” and study type “Expression profiling by array”. We identified and collected publicly available microarray data from 38 datasets with a total of 4,845 human kidney biopsy samples (Table 1). For each dataset, we carefully collected the platform information, annotations, and expression data. The gene expression measurements were made on 18 different platforms (Table 1). The commercial chips were designed by the manufacturers Affymetrix, Agilent and Illumina, and the custom chips were by Functional Genomics Facility at Stanford University and Bioinformatics and Gene Network Research Group at Zhejiang University. The number of probes varied widely from 1,347 to 54,675. In order to preserve as many genes as possible for further analysis we filtered out two datasets GSE26578 and GSE1563 with the lowest number of probes. We also filtered out 5 more datasets: GSE1743, GSE21785, GSE98320, GSE83486, and GSE109346, due to multiple absent or poor annotations such that it was impossible to confidently identify phenotypes associated with the individual samples. The dataset GSE343 was removed since the data of only log2 ratios of intensities in Cy5 and Cy3 channels were available. We found two datasets, GSE14700 and GSE82337, and one sample from GSE36059 with very sparse expression data and significant percentage of missing values, hence we filtered out these datasets as well. In the last step, we examined all datasets for any duplicate samples since many studies recycle previously published data. We were able to identify and remove 179 duplicate samples from GSE11166, GSE14328, GSE34437, GSE50058, and GSE72925 based on available metadata. We then used an R package DoppelgangR [23] that is based on calculations of correlations between samples and manually curated its results for any false positives and identified 257 samples as highly possible duplicates that were then also removed. Using the PCA plots, boxplots, and density plots, we analyzed the datasets for any presence of outliers. After stringent data quality control procedures, we composed the final dataset consisted of 28 studies with 2,273 samples. Their diagnostic annotations included 510 Acute rejection (AR) (including Antibody mediated rejection (ABMR), T cell mediated rejection (TCMR), AR, AR with Chronic allograft nephropathy (AR+CAN), Borderline rejection (BL), BL+CAN, Mixed rejection), 1,154 Stable (STA), and 609 Normal (i.e. biopsy taken prior to transplantation). The summary for the collected studies is represented in Table 1.

**Table 1.** Datasets collected from Gene Expression Omnibus (GEO). * Datasets not included into this study

| Study | PMID | Platform | Probes | Total | AR | STA | Normal | Year | Country |
| --- | --- | --- | --- | --- | --- | --- | --- | --- | --- |
| GSE343* | 12853585 | GPL271 LC-17 & GPL272 LC-20 | 28032 & 37632 | 41 | 26 | 0 | 15 | 2003 | USA |
| GSE1563* | 15307835 | GPL8300 [HG_U95Av2] Affymetrix Human Genome U95 Version 2 | 12625 | 31 | 7 | 10 | 9 | 2004 | USA |
| GSE1743* | 15476476 | GPL96 [HG-U133A] Affymetrix Human Genome U133A | 22283 | 41 | 0 | 41 | 0 | 2004 | USA |
| GSE7392 | 17397532 | GPL570 [HG-U133_Plus_2] Affymetrix Human Genome U133 Plus 2.0 | 54675 | 22 | 0 | 7 | 15 | 2007 | USA |
| GSE9493 | 19017305 | GPL570 [HG-U133_Plus_2] Affymetrix Human Genome U133 Plus 2.0 | 54675 | 56 | 21 | 21 | 14 | 2007 | Switzerland |
| GSE10419 | 20525436 | GPL887 Agilent-012097 Human 1A Microarray (V2) G4110B | 22153 | 29 | 0 | 29 | 0 | 2008 | Japan |
| GSE11166 | 19443638 | GPL570 [HG-U133_Plus_2] Affymetrix Human Genome U133 Plus 2.0 | 54675 | 49 | 1 | 27 | 21 | 2008 | USA |
| GSE14328 | 20150539 | GPL570 [HG-U133_Plus_2] Affymetrix Human Genome U133 Plus 2.0 | 54675 | 36 | 18 | 18 | 0 | 2009 | USA |
| GSE14700* | 20713790 | GPL8150 Stanford Microarray Functional Genomics Homo sapiens 34.6K | 34599 | 40 | 0 | 0 | 40 | 2009 | Austria |
| GSE21374 | 20501945 | GPL570 [HG-U133_Plus_2] Affymetrix Human Genome U133 Plus 2.0 | 54675 | 282 | 76 | 206 | 0 | 2010 | Canada |
| GSE22459 | 20813870 | GPL570 [HG-U133_Plus_2] Affymetrix Human Genome U133 Plus 2.0 | 54675 | 25 | 0 | 25 | 0 | 2010 | USA |
| GSE25902 | 21881554 | GPL570 [HG-U133_Plus_2] Affymetrix Human Genome U133 Plus 2.0 | 54675 | 88 | 27 | 37 | 24 | 2010 | USA |
| GSE26578* | - | GPL9301 Zhejiang university human 449 oligonucleotide array | 1347 | 85 | 34 | 51 | 0 | 2011 | China |
| GSE30718 | 22343120 | GPL570 [HG-U133_Plus_2] Affymetrix Human Genome U133 Plus 2.0 | 54675 | 19 | 0 | 11 | 8 | 2011 | Canada |
| GSE34437 | 23437201 | GPL570 [HG-U133_Plus_2] Affymetrix Human Genome U133 Plus 2.0 | 54675 | 36 | 17 | 7 | 12 | 2011 | USA |
| GSE34748 | 22335458 | GPL570 [HG-U133_Plus_2] Affymetrix Human Genome U133 Plus 2.0 | 54675 | 36 | 0 | 36 | 0 | 2011 | USA |
| GSE36059 | 23356949 | GPL570 [HG-U133_Plus_2] Affymetrix Human Genome U133 Plus 2.0 | 54675 | 410 | 122 | 280 | 8 | 2012 | Canada |
| GSE43974 | 25427168 | GPL10558 Illumina HumanHT-12 V4.0 expression beadchip | 47323 | 463 | 0 | 112 | 351 | 2013 | Netherlands |
| GSE44131 | 24030736 | GPL6244 [HuGene-1_0-st] Affymetrix Human Gene 1.0 ST Array | 29096 | 12 | 0 | 12 | 0 | 2013 | USA |
| GSE47097 | 23763497 | GPL6883 Illumina HumanRef-8 v3.0 expression beadchip | 47323 | 40 | 36 | 4 | 0 | 2013 | Netherlands |
| GSE48581 | 23915426 | GPL570 [HG-U133_Plus_2] Affymetrix Human Genome U133 Plus 2.0 | 54675 | 306 | 78 | 222 | 6 | 2013 | Canada |
| GSE50058 | 24127489 | GPL570 [HG-U133_Plus_2] Affymetrix Human Genome U133 Plus 2.0 | 54675 | 56 | 31 | 25 | 0 | 2013 | USA |
| GSE50084 | 26484130 | GPL6244 [HuGene-1_0-st] Affymetrix Human Gene 1.0 ST Array | 29096 | 48 | 28 | 20 | 0 | 2013 | USA |
| GSE52694 | 26176825 | GPL10558 Illumina HumanHT-12 V4.0 expression beadchip | 23719 | 14 | 0 | 14 | 0 | 2013 | Czech Republic |
| GSE53605 | 24698514 | GPL571 [HG-U133A_2] Affymetrix Human Genome U133A 2.0 | 22277 | 31 | 13 | 18 | 0 | 2013 | USA |
| GSE53769 | 20819195 | GPL16686 [HuGene-2_0-st] Affymetrix Human Gene 2.0 ST Array | 53617 | 28 | 0 | 10 | 18 | 2013 | Austria |
| GSE54888 | - | GPL6244 [HuGene-1_0-st] Affymetrix Human Gene 1.0 ST | 33252 | 54 | 0 | 0 | 54 | 2014 | Brazil |
| GSE57387 | 27452608 | GPL5175 [HuEx-1_0-st] Affymetrix Human Exon 1.0 ST | 17881 | 104 | 0 | 104 | 0 | 2014 | USA |
| GSE60807 | 26908771 | GPL6480 Agilent-014850 Whole Human Genome Microarray 4x44K G4112F | 41093 | 37 | 0 | 0 | 37 | 2014 | Austria |
| GSE65326 | 25307039 | GPL10558 Illumina HumanHT-12 V4.0 expression beadchip | 47323 | 6 | 0 | 6 | 0 | 2015 | Australia |
| GSE69677 | 27369853 | GPL14951 Illumina HumanHT-12 WG-DASL V4.0 R2 expression beadchip | 29360 | 76 | 24 | 0 | 52 | 2015 | Italy |
| GSE72925 | 27140517 | GPL570 [HG-U133_Plus_2] Affymetrix Human Genome U133 Plus 2.0 | 54675 | 46 | 10 | 36 | 0 | 2015 | USA |
| GSE76882 | 26990570 | GPL13158 [HT_HG-U133_Plus_PM] Affymetrix HT HG-U133+ PM | 17564 | 182 | 83 | 99 | 0 | 2016 | USA |
| GSE82337* | 27225518 | GPL10558 Illumina HumanHT-12 V4.0 expression beadchip | 47323 | 78 | 0 | 78 | 0 | 2016 | USA |
| GSE83486* | 28436117 | GPL10558 Illumina HumanHT-12 V4.0 expression beadchip | 47323 | 60 | ? | ? | 0 | 2016 | Norway |
| GSE98320* | 28614805 | GPL15207 [PrimeView] Affymetrix Human Gene Expression Array | 49395 | 1208 | ? | ? | 0 | 2017 | Canada |
| GSE109346* | - | GPL10558 Illumina HumanHT-12 V4.0 expression beadchip | 47323 | 26 | 0 | 26 | 0 | 2018 | Czech Republic |

### Data processing and normalization

Several pre-processing steps were applied prior to the main analysis. Raw fluorescence intensity data stored in .CEL or .txt files were downloaded and pre-processed depending on the platform. The data processing included background correction, log2 transformation, quantile normalization and probe to gene mapping using R language version 3.5.1 [24]. For the Affymetrix platform, we used the R package *SCAN.UPC* [25] available at Bioconductor [26] (http://www.bioconductor.org). In contrast to some other popular multi-array normalization algorithms like RMA [27] that estimate probe-level effects and standardize variances across arrays based on the information from a whole dataset, SCAN.UPC is a single-array method that normalizes every sample independently from other samples. This is considered as an advantage [25] since this approach is robust to any influence from possible outliers in the data. The database for the mapping between probes and Entrez gene IDs were taken from the BrainArray resource [28] version 22 (http://brainarray.mbni.med.umich.edu/Brainarray/Database/CustomCDF/22.0.0/entrezg.asp). For the Agilent and Illumina platforms, we downloaded non-normalized raw data and performed data processing using *neqc()* function within *limma* package [29] from Bioconductor. This algorithm [30] estimates parameters based on normal-exponential (normexp) convolutional model with joint likelihood estimation and with the help of negative control probes. The offset 16 was added to the intensities after the background adjustment by default, as it was shown as the most optimal value to improve FDR of the normexp algorithm [29]. However, some data from the Agilent and Stanford platforms did not contain any negative control probes. Therefore, we used similar processing steps manually to reproduce the methodology by applying *backgroundCorrect()* function from *limma* package by the *mle normexp* method with offset *16*. *Log2* transformation and quantile normalization was performed after this. The probe-gene mapping was implemented using the information from *biomaRt* database [31] or GPL files.

To gain additional statistical power from the large dataset we were able to collect, we chose to merge all the studies to perform a meta-analysis. In order to merge data from different studies and different platforms we had to first correct for potential batch effects. There are a number of papers that address this issue [32–36], but unfortunately, a one-fits-all solution doesn’t exist. Different normalization methods have their advantages and disadvantages [37–39] in removing batch effects, however, they can also become a critical problem in correcting the imbalanced data [32,36]. We examined the performance of several approaches that included *ComBat* [40] as a part of sva package [41], *Quantile Normalization (QN)* [42], *Remove Unwanted Variation (RUV)* [43] and *Harman* [44]. We identified superior batch effect minimization with *ComBat* in comparison to other methods and used it to normalize the data. A short description of our examinations can be found in *Supplemental materials*. The process of data aggregation, normalization and merging is schematically described in Fig 1a.

**Fig 1.**
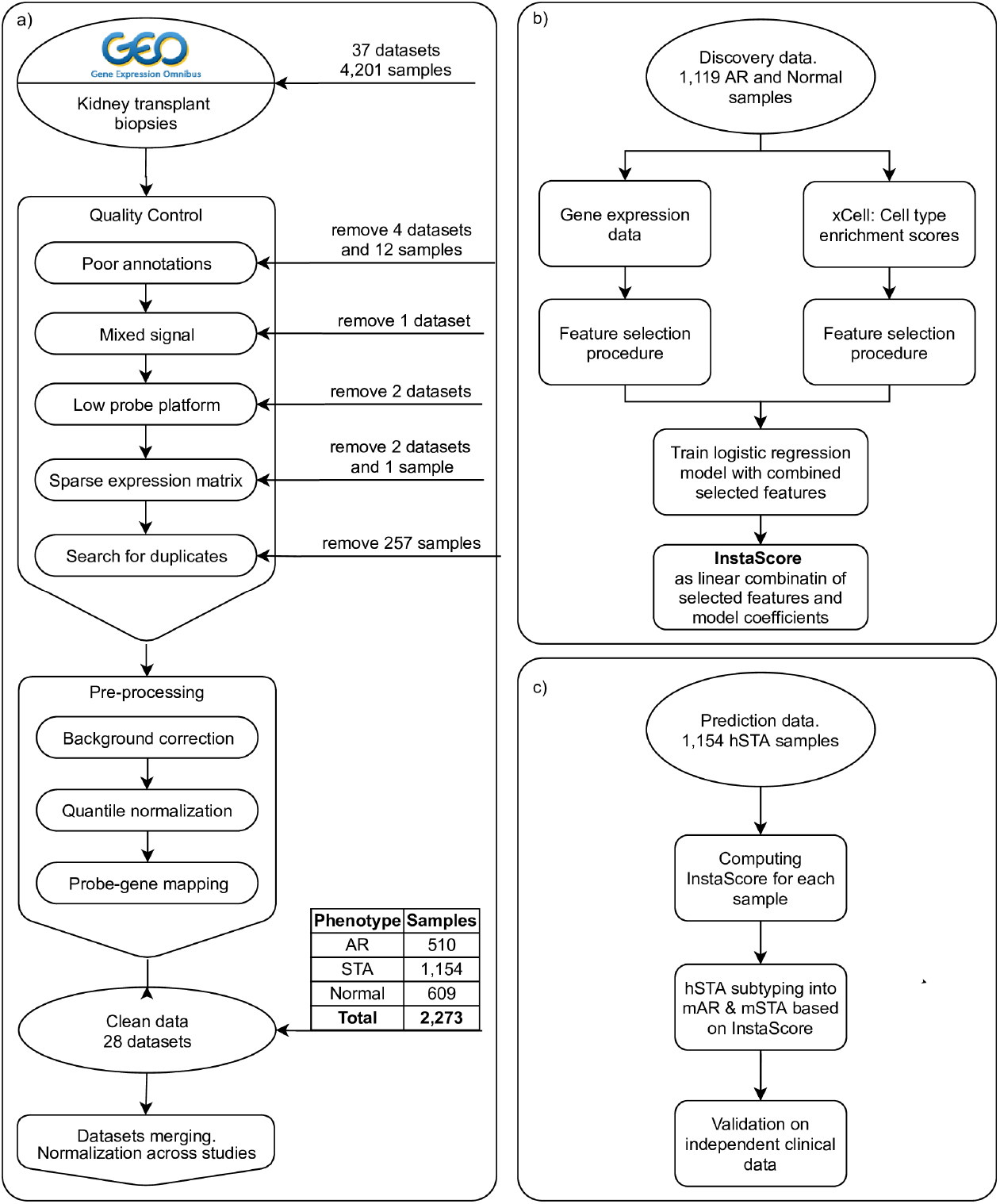
Flow chart. (a) Data pre-processing and Normalization. (b) Feature selection and creating the Instability Score (*InstaScore*). (c) Applying *InstaScore* to hSTA for sub-phenotyping and validating results.

### Differentially Expressed Gene (DEG) analysis

To identify differentially expressed genes in the first analysis AR vs Normal we used the Significance Analysis of Microarrays (SAM) [45] method that was implemented in the R package *siggenes* [46]. We utilized the false discovery rate (FDR) [47] with Benjamini-Hochberg procedure [48] for multiple testing correction and use the adjusted cutoff of 0.05. For the second level of significance, we selected only those genes that have the fold change greater than 1.5.

### Pathway Analysis

We leveraged the Gene Ontology database using the gene set enrichment analysis implemented in the R package *clusterProfiler* [49] to perform functional annotations for the significantly up- and down-regulated genes. We used FDR multiple correction method with the enrichment significance cut-off at level 0.05. For the gene network analysis, we utilized the STRING protein–protein association networks database [50] (https://string-db.org).

### Cell type enrichment analysis

In order to estimate the presence of certain cell types in biopsy samples, we leveraged a recently published cell type enrichment tool *xCell* [51]. xCell leverages gene expression data from microarray and RNA-seq experiments and is used to perform enrichment analysis for up to 64 immune and stromal cell types. We focused on 34 immune related and 11 non-immune cell types (Table S1) that we selected manually as relevant to the transplant injury process. In our analysis, we used a dedicated R package that is available on the author’s GitHub account for this purpose. This analytical method is a gene signatures-based method and converts the gene expression into cell type enrichment scores. The authors especially emphasize that this is not a deconvolution method that provides percentage of cell types containing in a tissue but rather the enrichment tool allowing to compare samples for each cell type but not otherwise. The enrichment scores for each cell type were used to compare AR and Normal samples and to identify cell types that are significantly different in individuals with AR as opposed to normal controls by performing the non-parametric two-sample Mann-Whitney-Wilcoxon statistical test. We utilized the multiple testing correction by using the Benjamini-Hochberg method. Adjusted p-value < 0.05 was used as the threshold.

### Feature selection procedure

In our efforts to select the most important features in distinguishing AR vs Normal samples, we performed the following steps. First, we split the whole data into training and testing sets in the ratio 80:20 and performed all feature selection procedures on the training set with benchmarking on the testing set. After identifying the significant features from a statistical test described above on the training set, we searched for features that correlate with the outcome no less than 2/3 of a maximum correlation value. In the final step, we applied Recursive Feature Elimination (RFE) technique with Random Forest (RF) model from the R package *caret* [52]. We used 5-fold cross validation (CV) technique with 100 repeats. To benchmark the model, we used the area under the ROC curve that is more suitable for data with some disbalance in outcome - in our case the ratio AR:Normal is 0.84. To decrease the bias of random split as well as to avoid the model overfitting, we also introduced the tolerance of 1% to the feature selection mechanism, i.e. the algorithm was choosing the simplest model with the smallest number of features that performs within range 99-100% of the best model.

To perform the steps described above, we adopted the R package *feseR* [53] and modified it to implement the parallel computations, the AUROC metric for model benchmarking, and the tolerance parameter of model performance. After the feature selection steps, we benchmarked the features with RF model on the testing set.

### Instability Score and hSTA sub-phenotyping

The method of sub-phenotyping hSTA samples is formed on creating a scoring system based on selected features and scoring the hSTA samples. Based on the score values, histologically STA samples are identified as *molecularly* AR or *molecularly* STA. We denoted this split as hSTA/mAR and hSTA/mSTA, respectively.

For our current analysis, two types of data were available: gene expression data and cell type enrichment scores (obtained computationally using xCell). Based on these two data types, we performed the feature selection procedure described above to find sets of genes and cell types that highly associated with AR. Next, we z-scaled each feature, a gene or a cell type, and built a logistic regression model with all features to identify feature importance as model coefficients. Using these coefficients, we created a linear formula to compute a score, that we called the *Instability Score* (or *InstaScore*):

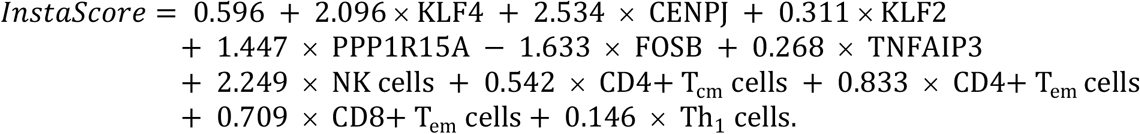

The positive *InstaScore* values separate AR from Normal samples which obtain negative values. By following the same steps, we then computed the *InstaScore* for the histologically STA samples and applied the zeroth threshold to obtain the split into mAR and mSTA subtypes. This whole approach is schematically represented in the form of the flow chart in the Figs 1b and 1c.

## Results

In the current study, we aimed to identify and characterize a phenotypic split of histologically STA (hSTA) kidney allografts that redefine inherent biological and functionally relevant differences that underlie the simple classification of a stable allograft by clinical function or histology alone, recognizing the redundancy of kidney reserve and the inherent discrepancies between different pathologists to interpret histology. In addition to better phenotyping hSTA allograft variabilities we also wanted to develop a robust, unbiased, quantitative classification method that could be integrated into clinical care, to better define immunologically quiescent renal allograft health. We leveraged all up-to date publicly available kidney biopsy microarray data and performed a feature selection procedure based on the Random Forest algorithm to identify a subset of genes and cell types that better distinguish AR and Normal samples. We then combined the feature selected genes and cell types into one score value, called *InstaScore*, applied it to hSTA samples, and identified the molecular subtype separation onto hSTA/mAR and hSTA/mSTA (Fig 1c). The prediction performance of the *InstaScore* was developed on public data and independently validated transcriptional data with linked clinical measurements.

### Datasets and Study Cohort

As a result of the data search in NCBI GEO and filtering procedures (see <u>Methods and Materials</u>) we were able to identify a robust dataset consisting of 2,273 samples including 510 AR, 1,154 STA, and 609 Normal samples from a total of 28 independent studies (Table 1). In the collection of the data on the Acute Rejection samples we included original annotations with investigator-based diagnoses of Antibody Mediated Rejection (ABMR), T-cell Mediated Rejection (TCMR), Acute Rejection with Chronic Allograft Nephropathy (AR+CAN), Borderline rejection (BL), BL+CAN, and Mixed rejection (TCMR and ABMR). Stable (STA) samples were histologically defined by each dataset as allografts not having rejection or any abnormalities including inflammation and fibrotic formations at the time of biopsy according to Banff’s classification criteria [15]. Normal samples were defined as donor engraftment biopsies at the time of transplantation.

### Differential Gene Expression Analysis Identifies Up-regulation of Immune Related Pathways in Rejection

Though prior gene expression studies have been done on understanding transcriptional perturbations in AR [54–57], leveraging thousands of kidney biopsy samples from public data gives us the statistical power to discover small biological effects during transplant rejection and injury and utilize this signature to unravel biological and molecular differences within hSTA transplants. In order to prepare all collected data in the form of separate studies, several preprocessing steps were carried out (see <u>Methods and Materials</u>). The scatter plots of the first two principal components of the gene expression data before and after the batch correction shows successful removal of batch effects (Fig S1). Next, we performed differential gene expression analysis AR vs Normal on the data (Benjamini-Hochberg adjusted p-value 0.05 and fold change 1.3) and identified 1,509 significantly differentially expressed genes including 848 up- and 661 down-regulated genes (Table S2). Further hierarchical clustering analysis on the significant genes based on Ward’s clustering technique was performed and showed the significant separation (p = 1.5E-14) between classes (Fig 2a). In addition, principal component analysis (PCA) confirmed the class separation (Fig S2). By leveraging the Gene Ontology database with the gene set enrichment analysis implemented in the R package *clusterProfiler* [49], we performed functional annotation for the significant genes and found, as expected, up-regulated genes were enriched in the regulation of the immune response, cell aggregation and activation, and innate immunity (Fig S3a). The down regulated genes were enriched in metabolic processes (Fig S3c). We also leveraged STRING, a tool for the functional protein association network analysis, [58] (https://string-db.org) for both up- and down-regulated genes showing significant connectivity between the sets of genes (Figs S3b, S3d).

**Fig 2.**
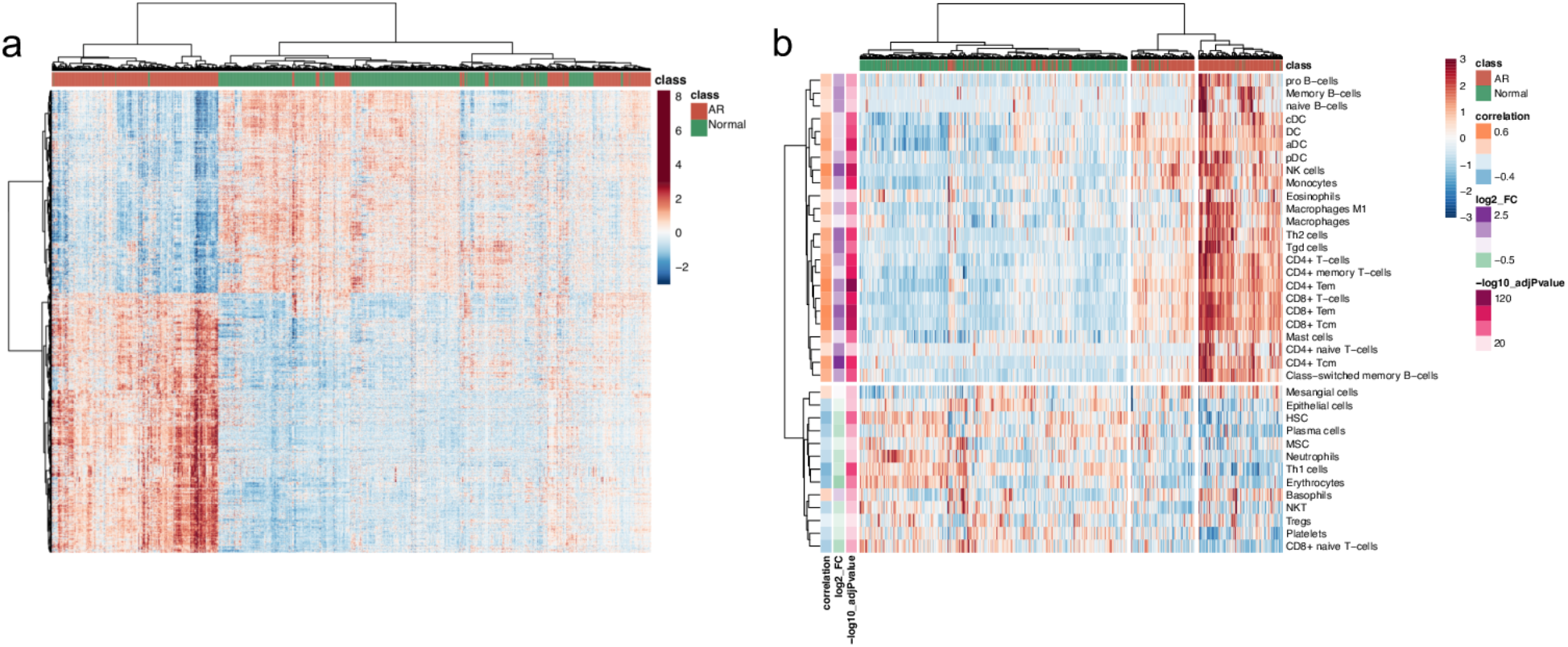
The heatmap plots for differentially expressed genes and significantly enriched cell types. (a) Heatmap clustering plot for significant genes from SAM analysis of AR vs Normals. (b) Heatmap clustering plots for significant cell types from the non-parametric Wilcoxon statistical test (BH p-adj < 0.05) in the analysis of AR vs Normals.

### Cell Type Enrichment Analysis Identifies Immune Cell Types Associated with Acute Rejection

To highlight the biological heterogeneity and to capture signals from infiltrating cell type specific effects in injured and stable kidney transplants, we performed cell type enrichment analysis that leverages cell type frequencies from the gene expression data. Currently, though single cell sequencing approaches are being developed, they are expensive, especially for large cohorts of this size, and often left over, or adequately preserved tissue is not available. Cell type deconvolution and enrichment methods provide an opportunity to look computationally at the cellular infiltration and activation and provide a unique assessment for STA allografts that may have subtle and sub-clinical rejection, not detectable by standard pathology. The xCell tool [51] computes enrichment scores for 64 immune and stroma cell types by leveraging tissue gene expression data. In our analysis, we focused on 45 cell types (Table S1) that would be more relevant for organ transplant. We performed non-parametric Wilcoxon tests to identify cell types that are significantly differentially enriched between AR and Normal samples. The p-values were adjusted to multiple testing correction with Benjamini-Hochberg method and with significant threshold of 0.05. We found 25 cell types (the majority of them are Lymphoid and Myeloid cells) that are significantly enriched in Acute Rejection, and 12 cell types (Immune, Stromal cells, and Hematopoietic stem cells) that are enriched in Normal kidneys. We show these results along with computed effect sizes and correlations with the case-control status in the form of the heatmap plot in the Fig 2b. On the heatmap, it is also interesting to see that AR samples fall into two main sub-clusters (p = 1.2e-10): one was mostly enriched in Lymphocytes, NK cells and Macrophages in contrast to another which had minimal Lymphocytes activation, and may represent temporal differences in rejection evolution or recovery. We observe that B cells, Dendritic cells, Macrophages, and T-cells form cell type specific sub-clusters that show the coordinated activation of the immune cells in kidney tissues. These results are in agreement with previous observations [54] that have shown AR sub-phenotypical splits by genes expression and cell types. Unsupervised clustering of hSTA along with AR and Normal samples revealed their heterogeneity hinting that some histologically stable samples have molecular signal closer to AR samples (Fig S4).

### Leveraging Feature Selection Procedure to Optimize AR classification

Following the feature selection procedure (Methods), we dramatically decrease the number of model features from all 1,509 differentially expressed genes only 6 pivotal upregulated genes: *KLF4, CENPJ, KLF2, PPP1R15A, FOSB, TNFAIP3* (AUC 0.98; Fig S5a), genes enriched as zinc finger proteins and expressed mostly in CD33+ Myeloid cells, and 5 cell types from the original set of 37 differentially enriched cell types: *CD4+ Tcm, CD4+ Tem, CD8+ Tem, NK cells,* and *Th1 cells*, with *CD4+ Tcm* having the largest effect size in this model (AUC 0.92; Fig S5b).

The feature selected cell types show a predominant role for infiltration and activation of effector T cells and NK cells in AR, and the feature selected genes appear to have broad cellular functions in AR, triggered by mononuclear activation and infiltration and collectively driving a variety of functions such as DNA recognition, RNA packaging, transcriptional activation and regulation of apoptosis. Interestingly, though previously the set of 11 genes in common rejection module (CRM) [9] identified from our cross organ (kidney, heart, liver, lung) meta-analysis study of transplant rejection is enriched in this current analysis, none of them make it to this final minimal feature selection set, suggesting this current 6 gene-set maybe more specific for absence of AR in the renal allograft, as the precise definition of a hSTA/mSTA allograft was not available in the earlier analysis.

A generated Random Forest classification model for these 6 genes and 5 cell types, internally validated using 5-fold cross-validation with 100 repeats, obtained an AUC of 0.98 (sensitivity 0.94, specificity 0.94) for the genes alone and an AUC of 0.92 (sensitivity 0.85, specificity 0.88) for the cell types for identification of a tissue sample with histologically confirmed AR (Fig 3a). We furthermore combined the feature selected genes and cell types into one score value, called *InstaScore* (Methods). By utilizing the *InstaScore*, we were able to perform the split into AR and Normal samples with slightly improved AUC 0.99 with sensitivity 0.95 and specificity 0.94 (Figs 3b, 3c).

**Fig 3.**
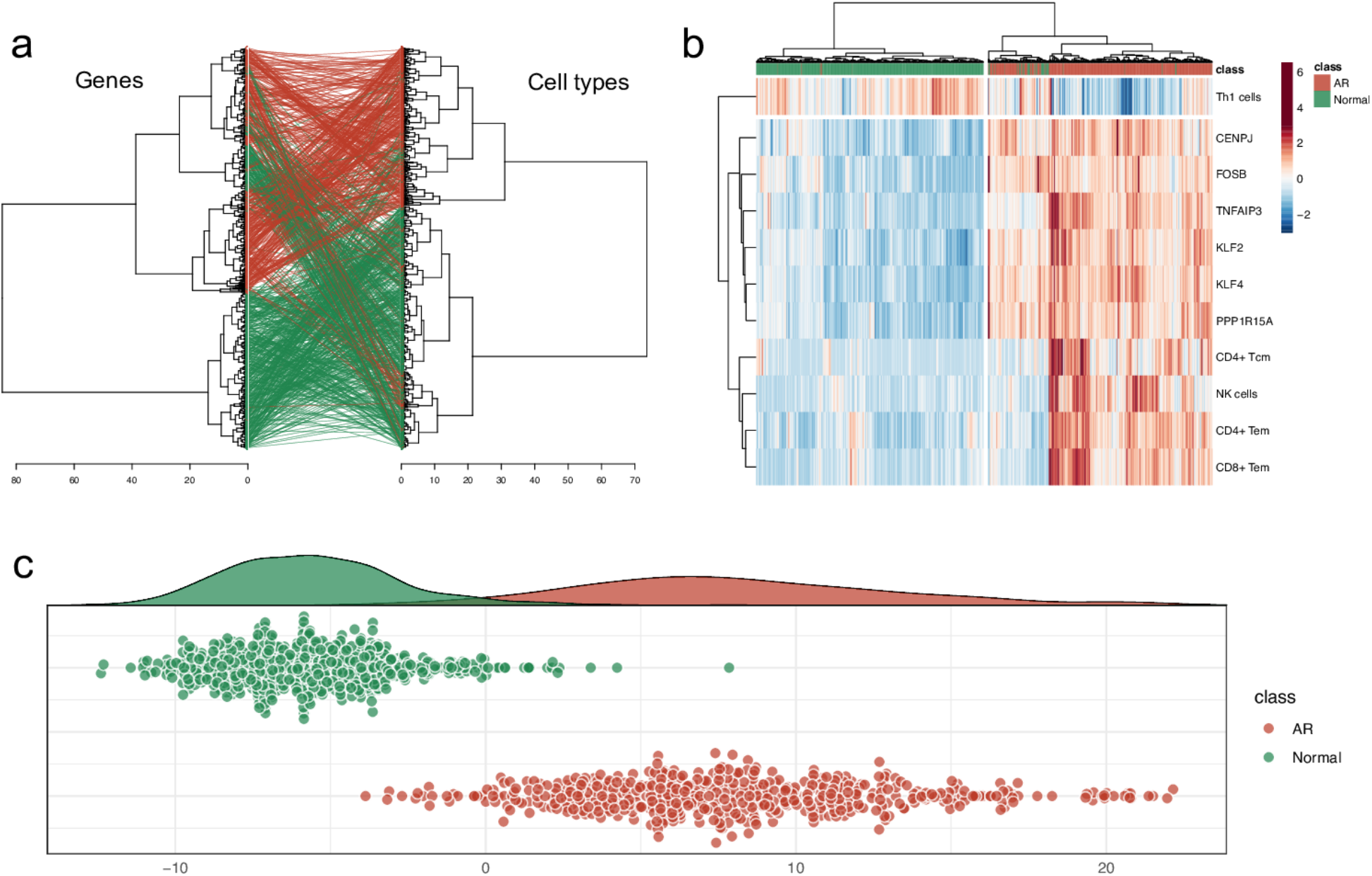
Feature selected genes and cell types and the Instability Score as their combination. (a) A comparison tanglegram between hierarchical clusterings based on Ward algorithm of AR and Normal samples for gene expression data with feature selected genes (left) and enrichment scores with feature selected cell types (right). The lines represent the tracking of each sample. Red color represents AR samples and green color represents Normal samples. (b) The heatmap plot of combined selected features with AR and Normal samples. (c) Instability Score plot for AR and Normal samples.

### Leveraging Selected Features to Create a Scoring Function to Carry out Precision Sub-phenotyping of Stable Samples

Having generated a powerful and robust tool of *in-silico* sample classification by the *InstaScore (Methods)*, we were able to apply this tool to the 1,154 transplant samples that were identified by pathologists/investigators in each of the datasets as *histologically* stable. With this approach, we are able recognize stable samples as more similar to Normal kidneys or as more similar to the rejected kidney allograft group, i.e. they would be *molecularly* stable (mSTA) or *molecularly* Acute Rejection (mAR). For this split of *histologically* stable transplants, we introduced new notations to reflect the deeper molecular sub-typing of functioning allografts: *hSTA/mSTA* for molecular and histological evidence of no rejection or tissue injury, and *hSTA/mAR* for allografts with clearer evidence of ongoing molecular rejection. The *InstaScore* identified 46% of all hSTA grafts (n= 528) as having mAR (Fig 4a). This “misclassification rate” of pathology is in a line with the previously reported discrepancies in transplant phenotyping across different pathologists [13,14].

**Fig 4.**
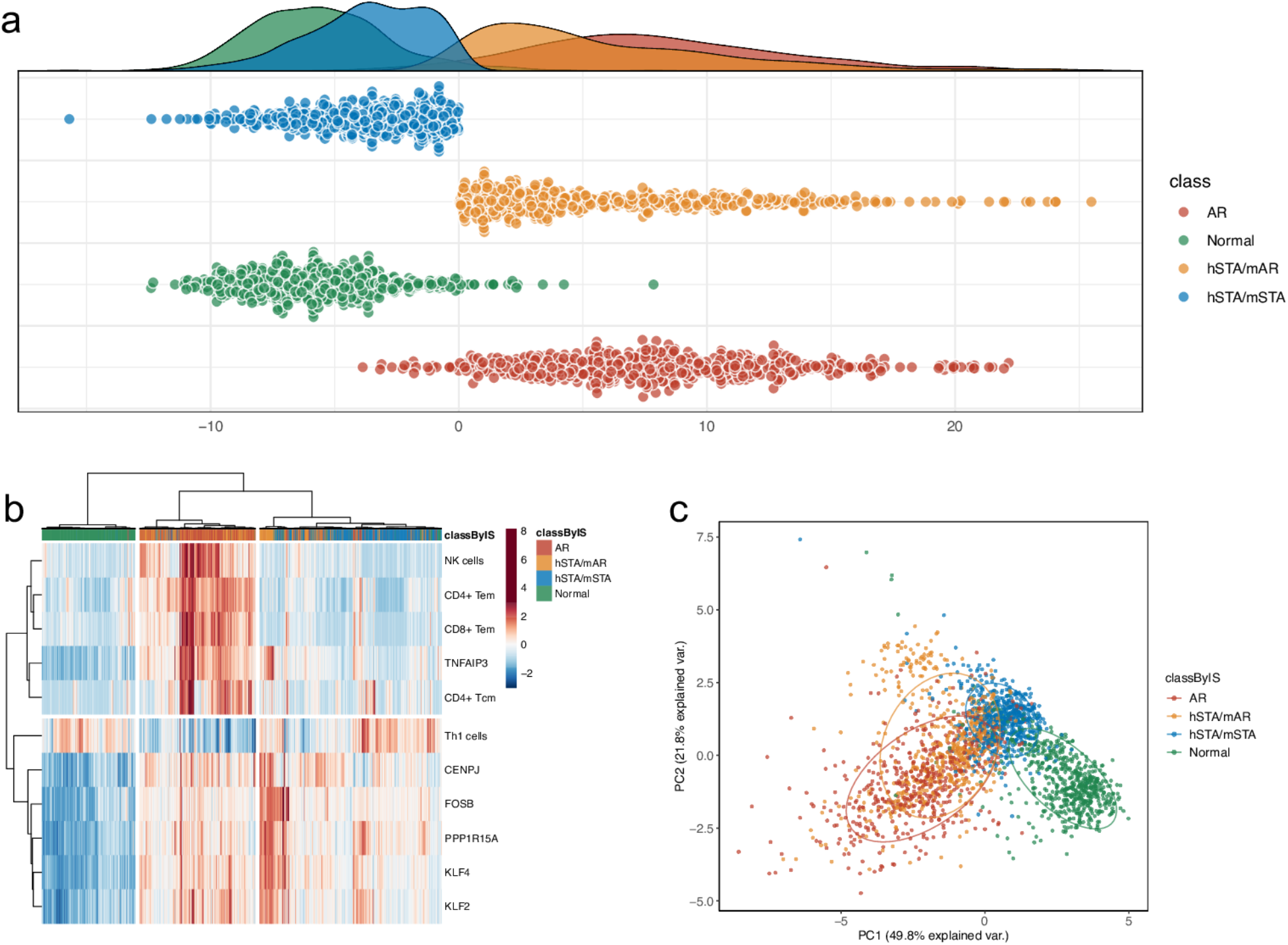
Plots of AR, sub-phenotyped hSTA, and Normal samples based on *InstaScore* results. (a) The Instability Score plots for AR, Normal, hSTA/mAR, and hSTA/mSTA samples. (b) Heatmap and (c) PCA plots of AR and Normal samples with sub-phenotyped hSTA samples based on the *InstaScore*.

We represented the scores for each sample in the form of the scatter plot (Figs 3c,4a). The *InstaScore* can significantly distinguish AR and Normal samples (p = 7e-179; Fig 3c), giving the ability to identify hSTA/mAR and hSTA/mSTA samples (p = 1e-188; Fig 4a) by thresholding with 0. From visualizing all the data with the selected features in the form of the heatmap (Fig 4b) and PCA plot (Fig 4c) and coloring sub-phenotypes based on the *InstaScore* one can notice that hSTA/mAR samples clustered nearly perfectly with AR samples separately from hSTA/mSTA samples and could be seen as ones in the intermediate stage between Normal and AR samples.

### Validation of hSTA sub-phenotyping using clinical follow-up data

In order to further demonstrate the functional relevance of the *InstaScore* by gene expression and cell types, we explored the clinical utility of the *InstaScore* in an independent microarray data set from 67 unique STA patients (stable clinical graft function, no DSA, no AR) from a randomized controlled clinical trial (NCT00141037) with transcriptional data on serial protocol kidney transplant biopsies at 0, 3, 6, 12, and 24 months [59,60] and with longitudinal functional outcomes up to 5 years after initial engraftment. We had the unique opportunity to test the impact of the locked *InstaScore* on the change of estimated glomerular filtration rate (eGFR) and graft loss events over this time period. We correlated the *InstaScore* values with those outcomes and found higher correlation values for cell type infiltration/activation model with delta eGFR (Fig 5a) r = 0.52 (p = 6.4e-6) and graft loss events r = 0.17.

**Fig 5.**
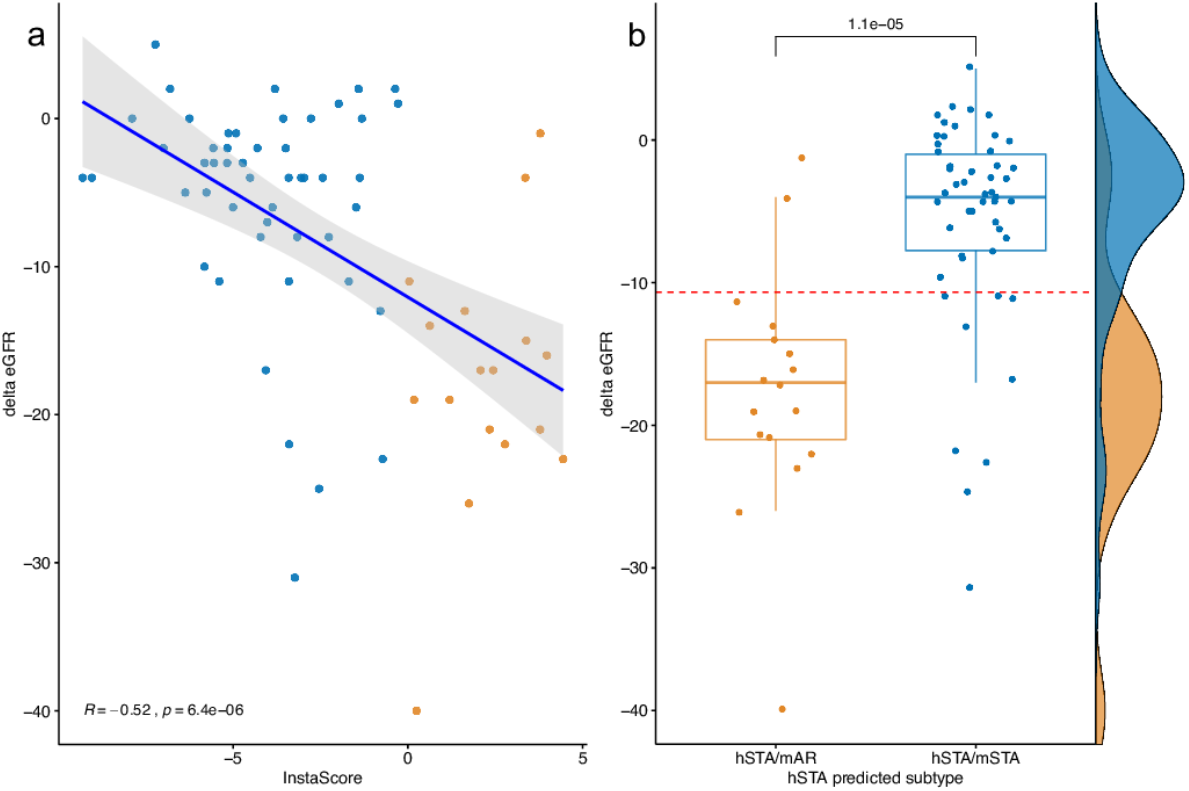
Validation plots on the independent clinical data. (a) Correlation plot between change in eGFR over 1-5 year period after biopsy and the *InstaScore* (cor = −0.52, p = 6.4e-6). (b) Boxplot of delta eGFR distributions for predicted hSTA subpopulations by the *InstaScore* (p = 1.1e-5).

Based on a separation of the hSTA groups by the *InstaScore*, the predicted hSTA sub-phenotypes, showed a significant separation over the 5-year follow-up with a delta eGFR separation of greater than −10. Fig 5b represents boxplot comparisons of *InstaScores* between predicted sub-phenotypes with this delta eGFR thresholding of >−10 and shows that the groups difference was significant (p = 1.1e-5). Therefore, we found that the *InstaScore* on the 6 month protocol biopsy can be valuable for predicting graft function outcomes 5 years later and thus can help differentiate those patients who are more likely to have progressive graft injury and decline in graft function over time, even though the 6 month protocol biopsy or DSAs at the 6 month time point post-transplant would not have been able to provide the same predictive discriminatory power.

## Discussion

In this study we identified discrepancies in the current histological recognition of stable allografts and demonstrated a new Instability Score (or *InstaScore*), a linear combination of selected features based on the input from tissue gene expression and inferred cell type biopsy molecular data modeled on AR biology, that provides precise (sub-)phenotyping and early recognition of molecular and cellular rejection of otherwise functionally and histologically stable allografts on protocol biopsies. Unlike other published studies by others [61–65] and our group [2,54,66,67] that have only studied gene expression transcriptional perturbation in AR, this is the first development and application of a combined genes and cell types into Instability Score in tissue that can rapidly reclassify samples into molecularly and even more importantly, functionally relevant, diverse groups: those most similar to the normal kidneys (hSTA/mSTA) and those most similar to the rejected kidneys (hSTA/mAR).

We employed machine learning methods to better understand the underlying mechanisms in stable kidney transplants and show possible reasons for discrepancy in histological diagnostics. By taking advantage of the statistical power of a large collective gene expression dataset and adopting a novel computational tool of cell type enrichment analysis, xCell [51], we refined the molecular and cellular profile of AR to narrow the signature to specific genes and cell types that precisely map rejection, and then used the logistic regression algorithm to build a linear combination of features, the *InstaScore*, to identify two major subgroups of samples, otherwise described homogenously as *stable* in the public datasets. About half (54%) of the histologically defined STA (hSTA) had immune quiescence and molecular similarities to the healthy donor kidney, and have been redefined in this study as histologically and molecularly stable (hSTA/mSTA), a large number (46%) of the remaining biopsies had molecular evidence of rejection (hSTA/mAR) and should not have been classified as stable allografts. The distribution of the hSTA/mAR stable biopsies were found to be scattered across multiple datasets, supporting that their presence is not due to failure of histological characterization at any particular transplant program. Independent validation of the *InstaScore* was conducted on microarray data from a set of biopsies not utilized for model training and identified subgroups of hSTA/mAR phenotypes again on 6 months protocol biopsies, from otherwise presumed functionally and histologically stable transplant recipients. The *InstaScore* application on these biopsies, where long term outcome data was available, stratified these kidney transplant recipients as being at greater risk of chronic graft rejection, more progressive interstitial fibrosis and tubular atrophy and thus, greater loss of graft function over subsequent follow-up. This critical information, if available to the physician in real time, can allow for recipient immunosuppression stratification, aggressive clinical follow-up and graft salvage; all of which are critical unmet needs in transplantation.

The 6 feature selected genes, *KLF4, CENPJ, KLF2, PPP1R15A, FOSB*, and *TNFAIP3*, contributed into the *InstaScore*, are significantly dysregulated and were highly associated with AR in comparison to normal kidneys. As expected, these significantly upregulated genes are enriched in the regulation of the immune response, cell aggregation and activation, and innate immunity. *KLF2, KLF4*, and *TNFAIP3* belong to zinc finger superfamily, where *KLF 4* and *2*, both belongs to the same group of Kruppel (zinc-finger) family of transcription factors and are known to regulate kidney injury/disease [68]. *KLF2* is vasoprotective and *KLF4* is renoprotective, both are enriched in the endothelium and have overlapping functions and similar downstream targets [69–71]. In AR, increased expression of *KLF 4* and *2* is also associated with endothelial ischemia reperfusion injury [72]. Zinc finger protein *TNFAIP3* has anti-apoptotic and anti-inflammatory functions and its expression localizes mainly to endothelial cells and infiltrating myeloid cells and expression of TNFAIP3 by infiltrating T cells (myeloid cells) results in adverse clinical outcomes[73–75]. *CENPJ* functions as a transcriptional coactivator in the STAT5 signaling pathway, and TNF-induced NF-kappaB-mediated transcription [76,77], both central regulators of inflammation. Phosphatase *PPP1R15A* is only expressed in stressed cells and negatively regulate acute kidney injury vis Type 1 interferon (IFN). Its overexpression has been observed in chronic inflammatory diseases such as lupus, which also has a type I IFN signature [78]. Type 1 IFN has a direct effect on CD8+ T cells, promoting clonal expansion and memory T-cell, plasma cell differentiation, and enhance B-cell responses [79]. *FOSB* expression has been associated with the progression of renal disease including IgA nephropathy, leading to end-stage renal failure [80]. Thus the discovery gene signatures from kidney biopsies in the *InstaScore* are crucial for - endothelial cell integrity, T-cell expansion and activity, and have been previously associated with inferior graft function and rejection [81,82]. Importantly, the utility of this gene set in the *InstaScore* prediction is significant, so application of a rapid PCR based assay on formalin fixed paraffin embedded slices from the pathology samples, could allow for triaging the molecular burden of injury on a protocol biopsy, without the need to obtain an additional “research” core for snap freezing or RNAlater, is generally recommended by previous studies. This approach has been developed and published recently by our group [83,84].

The 5 feature selected cell types are also in line with organ rejection biology[85]. AR is associated with early infiltration of CD8+ T and CD4+ T cells, macrophages, NK cells, and B cells [86,87]. Transplanted kidneys lose their function due to both immunologic and metabolic mechanisms but the most predominant cause of graft loss, according to the latest Banff classification of renal allograft pathology, is alloimmune response with the major role of T-cell in acute rejection [88]. From our feature selection procedure, we identified 5 cell types from the original set of 64 cell types (from which 37 were enriched in AR): CD4+ Tcm, CD4+ Tem, CD8+ Tem, NK cells, and Th1 cells, with CD4+ Tcm having the largest effect size. NK cell are pathogenesis-linked markers for antibody mediated rejection [89] and are known to regulate diverse T cell responses promoting T-cell proliferation and graft rejection [90].In the immunologic response to the allograft, T cells terminally differentiate and divide into central memory T cells (Tcm), and effector memory T cells (Tem) [91]. Tem can be both CD8+ and CD4+, where both produce IFN-gamma, IL-4, and IL-5 within hours following antigenic stimulation, however, CD8+ Tem carry large amounts of cytotoxic molecules like granzyme, granulysin and perforin [92,93]. In our feature selection procedure, we identified both, CD4+ Tem, CD8+ Tem, as the two enriched cell types for AR, but the classification model recognized CD4+ Tcm having the largest effect size in this model. This is likely because T central memory cells mediate reactive memory, which has higher sensitivity to antigenic stimulation, is less dependent on co-stimulation, and provides more effective stimulatory feedback to dendritic cells (DC) and B cells. Compared to Tem, Tcm are characterized by slow effector function, with expansion and differentiation of T cells in response to repeat antigenic stimulation [94]. Thus the higher enrichment of CD4+ Tcm in the *InstaScore* may suggest the inherent, pre-determined ability of a hSTA/mAR graft, in the absence of substantive active inflammation at the time of biopsy, to be primed to differentiate into Tem even with lower levels of subsequent antigen recognition, such as with varying exposure to immunosuppression[95]. Thus, the *InstaScore* may be identifying a primed hSTA graft that is greater risk to subsequent immune injury and allograft damage as it already carries a primed signature of mAR, than an allograft that is classically stable (hSTA/mSTA).

Conversely, Th1 cells are less enriched in the *InstaScore*, which is a little surprising as Th1-cell subsets are pro-inflammatory, the Th2-cell subsets anti-inflammatory, and Th1–Th2 balance is regarded as the major mechanism of rejection [96]. The ratio Th1/Th2 in the *InstaScore* was decreased in comparison to Normal healthy control samples by 4.6 times, although an increase in the ratio is associated with AR as observed before [97,98]. We believe this difference may relate to the temporal nature of the TH1/Th2 state and a skew towards more M2 macrophages rather than M1 macrophages, as we are really detecting sub-clinical molecular rejection, rather than full-blown, clinically relevant AR, where the reverse findings are usually observed. Increased numbers of M2 than M1 macrophages have been found in some studies to correlate with graft-loss [99]. KLF4 expression in increased in the *InstaScore*, and interestingly this gene plays a key role to polarize macrophage into the M2 subset [100], which undertake host defense and wound healing/tissue remodeling tasks[101] and thus can be found to be enriched in renal fibrosis [102].In the AR bopsies in this study, as expected, M1 macrophage signatures were higher in AR [103]. Thus, the macrophage subsets show a dynamic state of skewing the preponderance of M2 to M1 macrophages in the process of immunological priming from pre-AR and full blown AR, and the *InstaScore* uncovers the state of AR priming or pre-AR in histologically and clinically stable kidney transplants, further highlighting the value of identifying hSTA grafts that are thus at greater risk of AR over time.

Given the design of the study, there are a few inherent limitations. First, this study was based on publicly available data with limited access to clinical and demographic reports in each of the public data. We excluded 5 datasets and 12 samples from the study as they lacked any detailed annotations. Secondly, we found 257 duplicates where some had slightly different histological annotations based on study aims, thus we had to carefully filter samples to minimize errors in sample identification. Thirdly, as the gene expressions in some datasets used in the study were measured using much older technologies, we excluded two sets of data from the current analysis. The presence of batch effects was recognized issue in this kind of analysis and can be challenging to control for as it can contribute to false positive findings in meta-analysis studies, but we believe that the testing of multiple normalizing methods provides a robust methodology for greater confidence in the data.

In conclusion, we have generated an Instability Score for histologically and clinically stable allograft profiled by microarray or RNASeq technologies (*InstaScore*) - the linear combination of feature selected 6 genes and 5 cell type. We have demonstrated that when we apply this Score to 1,154 transplant samples that were identified by pathologists as hSTA samples it segregates hSTA samples, that would otherwise be grouped into a single bucket, now into quite different molecular subtypes, that were subsequently confirmed by independent data analysis to be very functionally diverse over time. Thus, thus is the first study that highlights that even among histologically stable kidney transplants there is wide heterogeneity that has to be addressed in transplant diagnostics and analysis. There is a need of a better and more precise method that can sub-phenotype stable allografts into those that are stable at all biological levels and those that are slowly sliding towards rejection. Based on the application of the *InstaScore* for transcriptional data, immunosuppression adjustments could be made in patients to preserve allograft function. We uncovered that accepting all samples classified as “stable” from different investigators and datasets will neglect the important biological diversity that naturally exists in the histological definition of a “stable” sample and will necessarily lead to false positive or false negative discoveries when attempting to study and fully understand allograft rejection.

## Acknowledgments

We would like to thank Tara Sidgel, Jul Liberto, and Parhom Towfighi for collecting and organizing clinical data, to Silvia Pineda and Jieming Chen for fruitful discussions and suggestions, and all members of the SarwalLab and SirotaLab who supported all study efforts. We are also thankful to patients and their families whose deidentified samples have provided the needed data to make these study findings.

## Supplementary Figures

**Fig S1.**
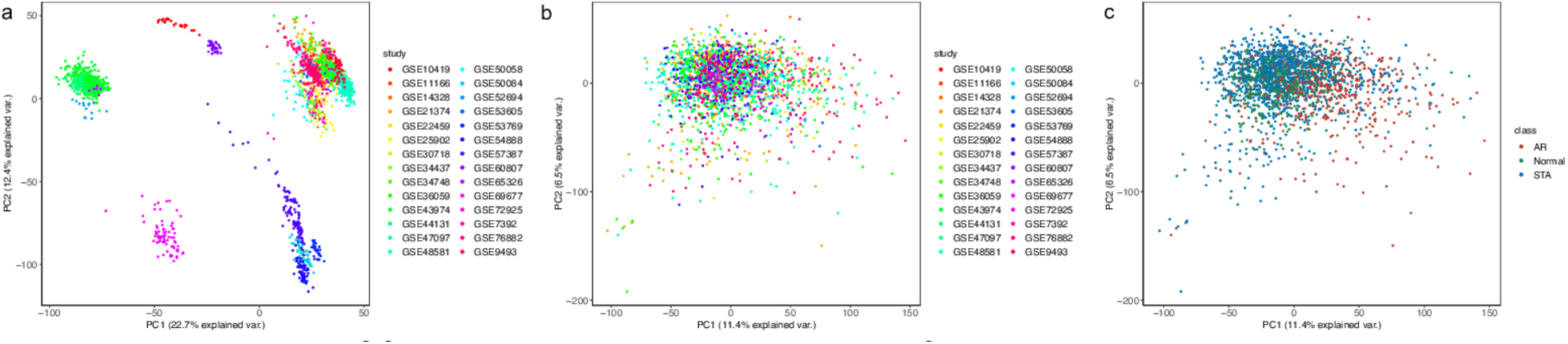
Scatter plots of first two principal components of gene expression data. PCA plots (a) before normalization colored by study, (b) after normalization with ComBat colored by study, (c) after normalization colored by phenotypes.

**Fig S2.**
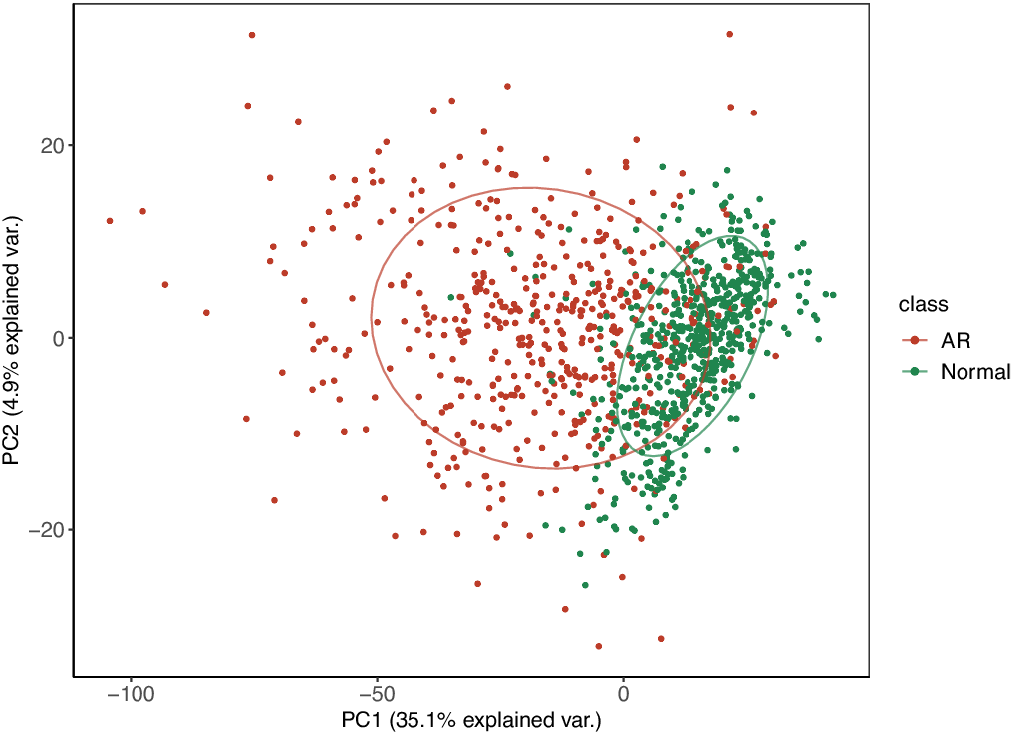
PCA clustering plot for differentially expressed genes from analysis of AR vs Normals.

**Fig S3.**
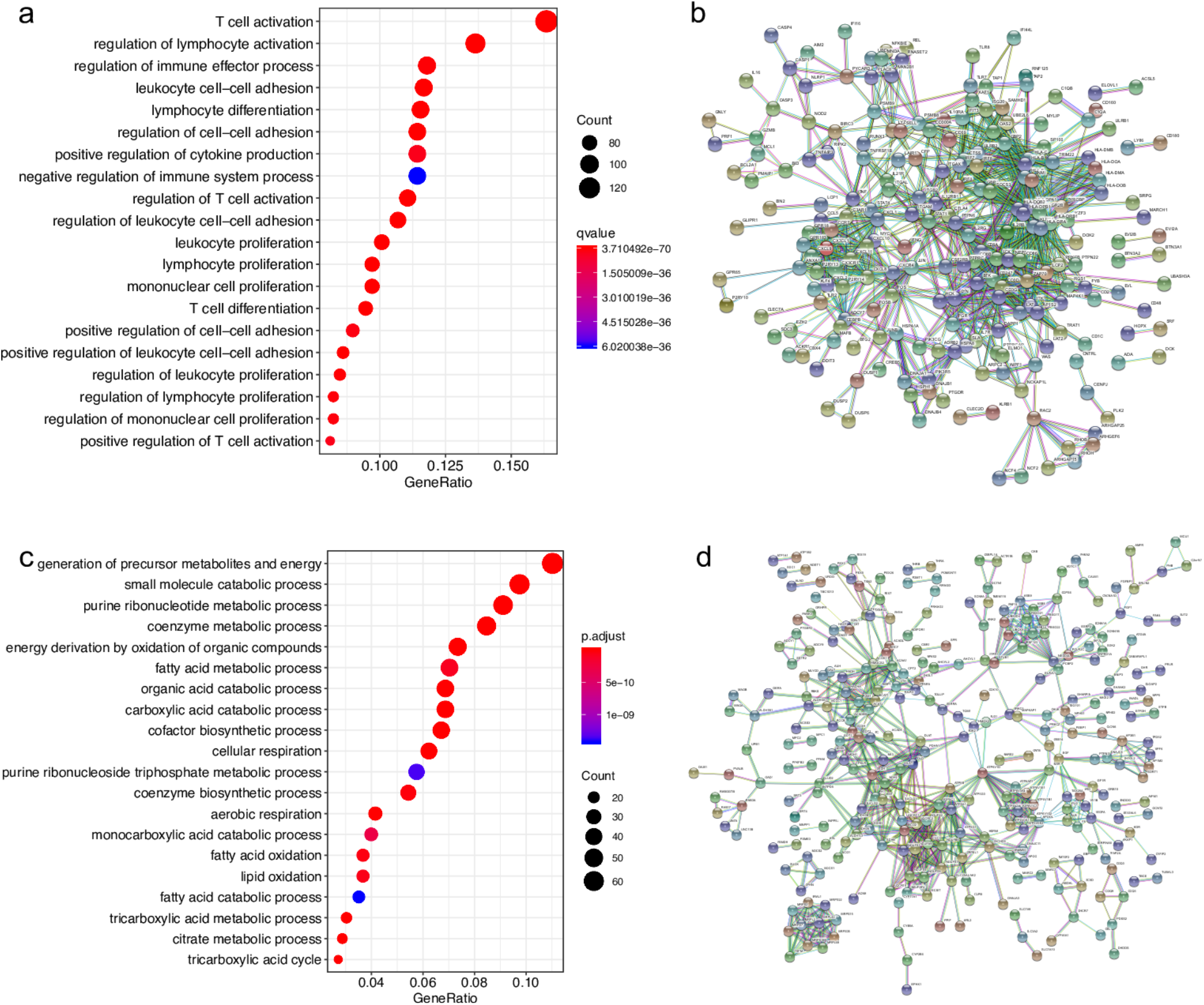
Pathway enrichment analysis of DE genes. The top 20 GO terms enriched among (a) up- and (c) down-regulated genes in AR vs Normal analysis. Functional networking with STRING for the differentially expressed (b) up- and (d) down-regulated genes. We used the minimum required interaction score equal to 0.9. The edges mean a type of interaction evidence. The colors mean as follows. The known interactions are cyan for curated databases and magenta for experimentally determined interactions. The predicted interactions are represented in green for gene neighborhood, red for gene fusions, and blue for gene co-occurrence. The other types of interactions are in lime based on text mining, black on co-expression, and indigo on protein homology.

**Fig S4.**
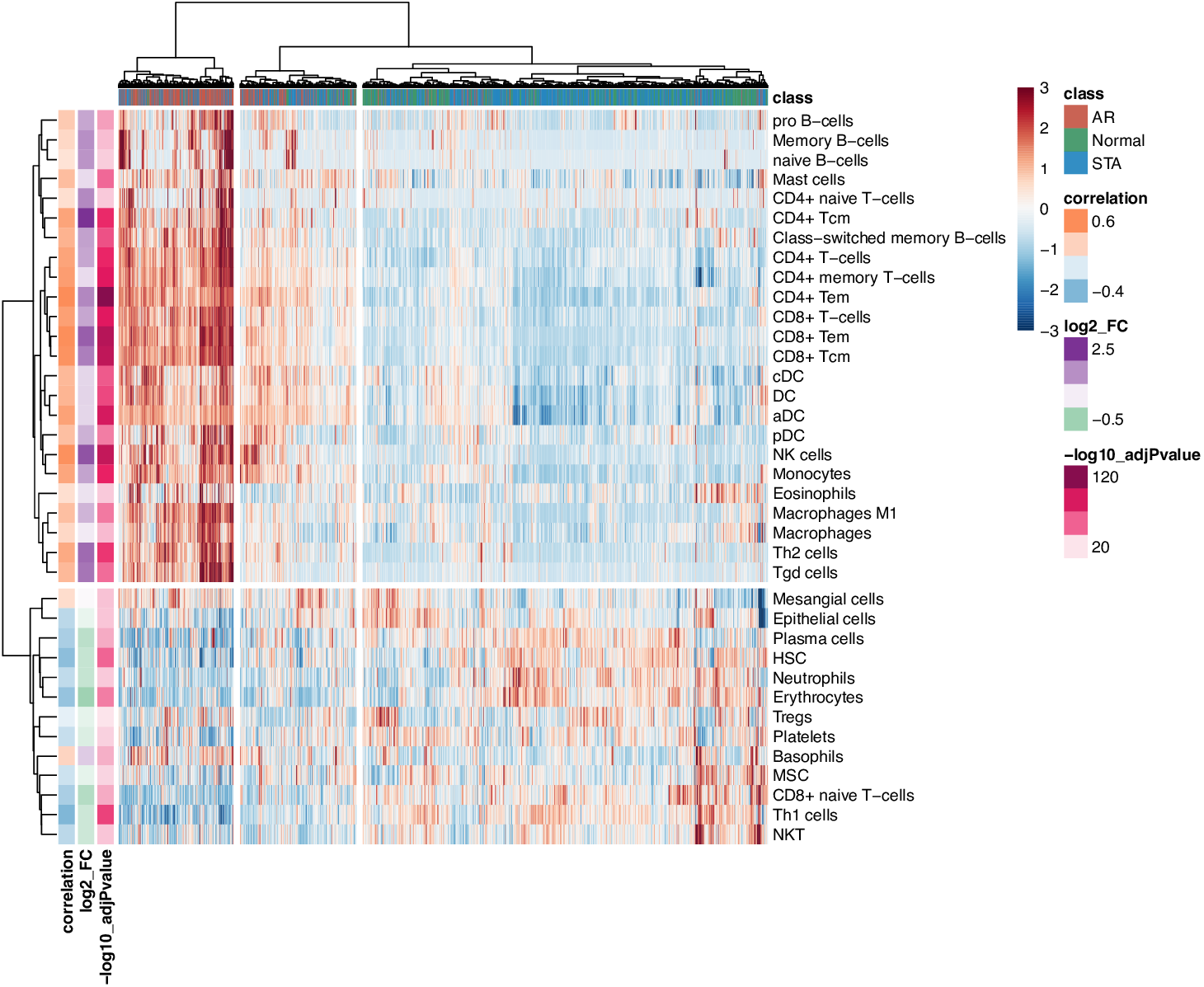
The heatmap of enrichment scores of significant cell types from the AR vs Normal comparison. The colored vertical bars represent results from AR vs Normal analysis. The plot combines all AR, hSTA, and Normal samples and shows clustering of some hSTA samples together with AR hinting to possible hidden inflammation processes going in those grafts.

**Fig S5.**
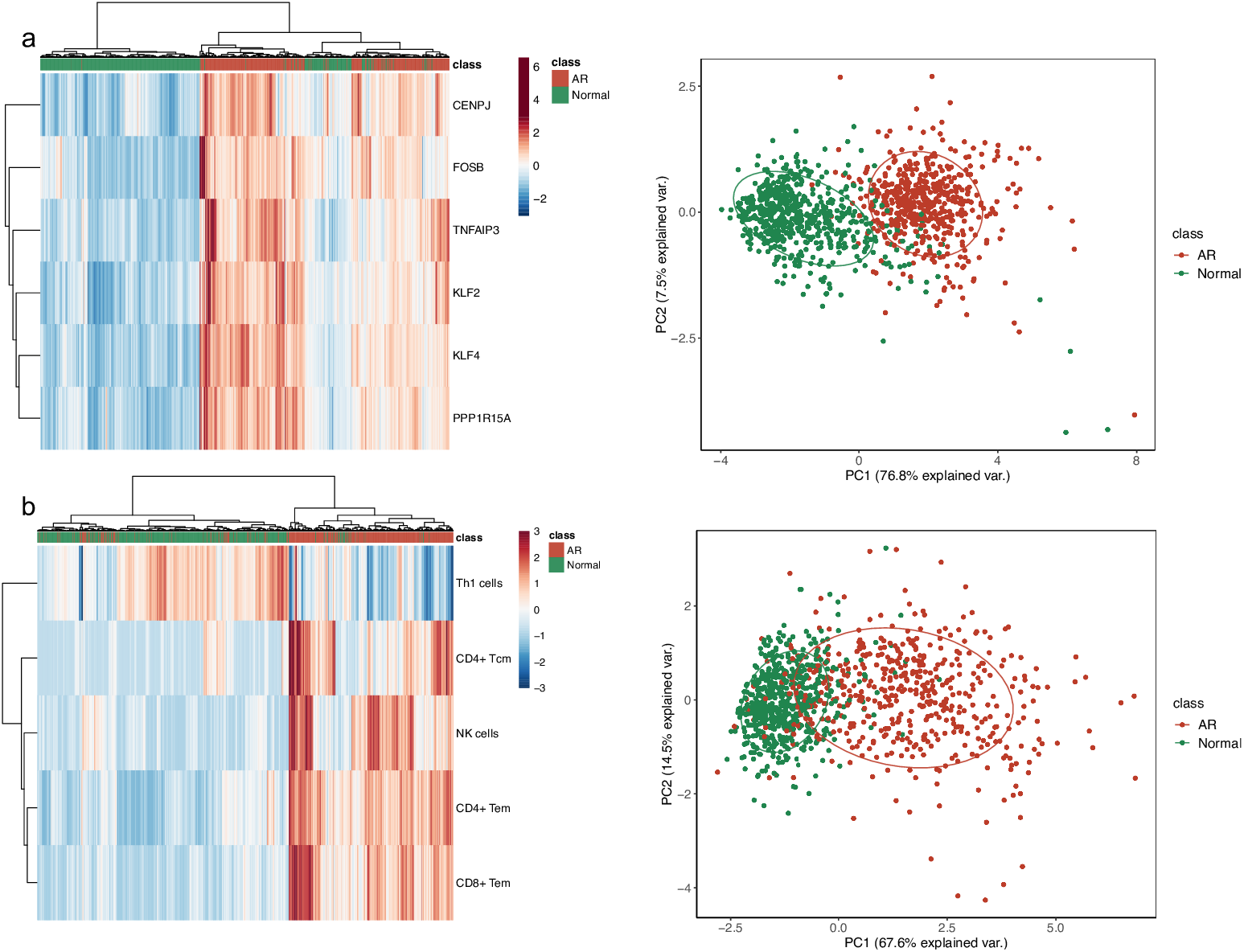
Plots of feature selected genes and cell types for all AR and Normal samples. (a) Heatmap and PCA plot of feature selected gene expression. (b) Heatmap and PCA plot of feature selected cell types.

**Fig S6.**
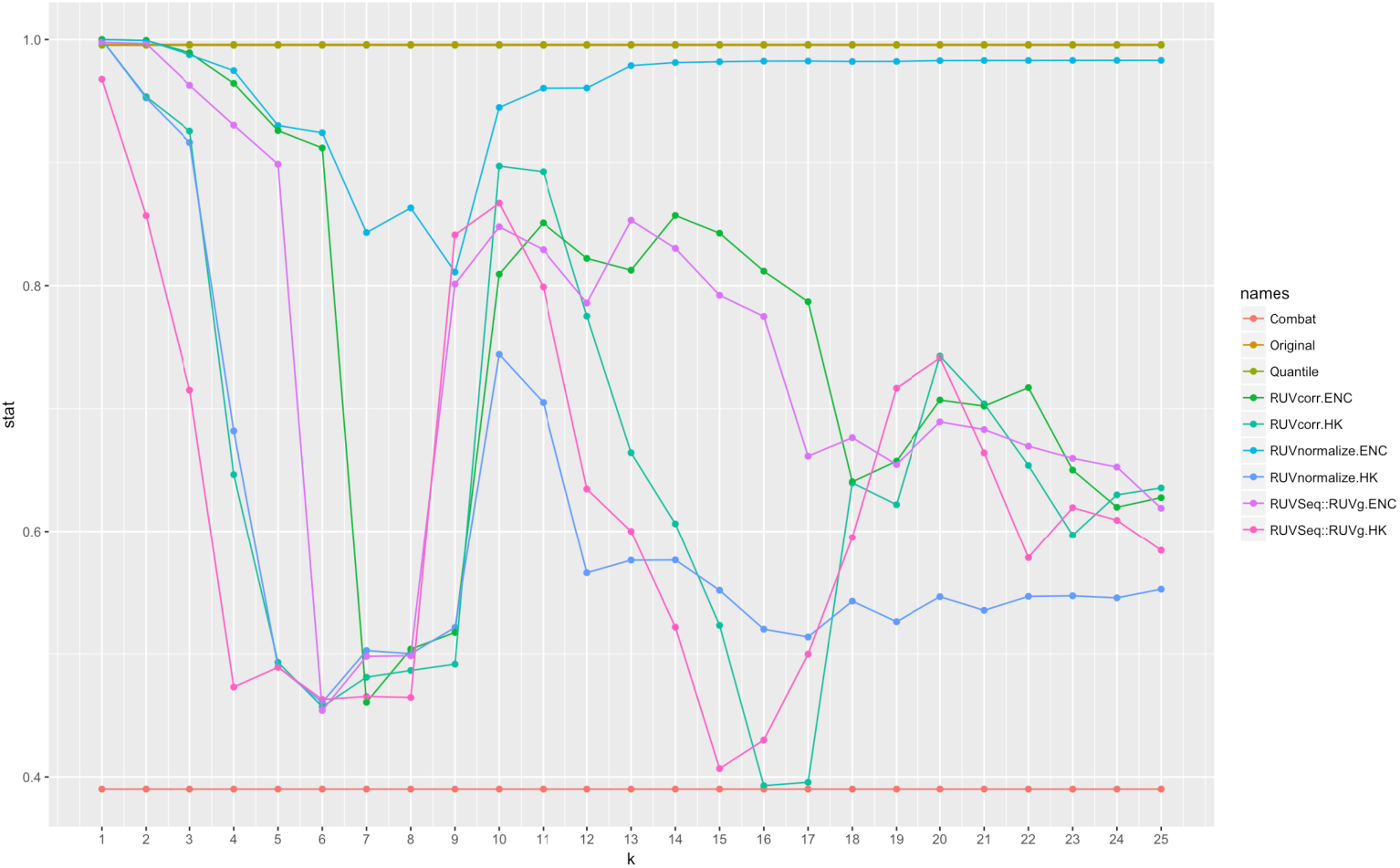
The combined banchmark based on p-value, delta statistic and the percentage of variability for batch correction methods tested. HK – Housekeeping genes, ENC – Empirical Negative Control genes

## Supplementary Table

**Table S1.** Cell types used in cell type enrichment analysis with xCell.

| Cells | Family | Type |
| --- | --- | --- |
| Erythrocytes | HSC | HSC |
| HSC | HSC | HSC |
| Megakaryocytes | HSC | HSC |
| Platelets | HSC | HSC |
| B-cells | Immune | Lymphoid |
| CD4+ memory T-cells | Immune | Lymphoid |
| CD4+ naive T-cells | Immune | Lymphoid |
| CD4+ T-cells | Immune | Lymphoid |
| CD4+ Tcm | Immune | Lymphoid |
| CD4+ Tem | Immune | Lymphoid |
| CD8+ naive T-cells | Immune | Lymphoid |
| CD8+ T-cells | Immune | Lymphoid |
| CD8+ Tcm | Immune | Lymphoid |
| CD8+ Tem | Immune | Lymphoid |
| Class-switched memory B-cells | Immune | Lymphoid |
| Memory B-cells | Immune | Lymphoid |
| naive B-cells | Immune | Lymphoid |
| NK cells | Immune | Lymphoid |
| NKT | Immune | Lymphoid |
| Plasma cells | Immune | Lymphoid |
| pro B-cells | Immune | Lymphoid |
| Tgd cells | Immune | Lymphoid |
| Th1 cells | Immune | Lymphoid |
| Th2 cells | Immune | Lymphoid |
| Tregs | Immune | Lymphoid |
| aDC | Immune | Myeloid |
| Basophils | Immune | Myeloid |
| cDC | Immune | Myeloid |
| DC | Immune | Myeloid |
| Eosinophils | Immune | Myeloid |
| iDC | Immune | Myeloid |
| Macrophages | Immune | Myeloid |
| Macrophages M1 | Immune | Myeloid |
| Macrophages M2 | Immune | Myeloid |
| Mast cells | Immune | Myeloid |
| Monocytes | Immune | Myeloid |
| Neutrophils | Immune | Myeloid |
| pDC | Immune | Myeloid |
| Epithelial cells | Non-Hematopoietic | Epithelial |
| Endothelial cells | Non-Hematopoietic | Stroma |
| Fibroblasts | Non-Hematopoietic | Stroma |
| ly Endothelial cells | Non-Hematopoietic | Stroma |
| Mesangial cells | Non-Hematopoietic | Stroma |
| MSC | Non-Hematopoietic | Stroma |
| mv Endothelial cells | Non-Hematopoietic | Stroma |

## Supplementary methods

### Cross-study normalization in data merging

We have examined several batch correction methods: *ComBat* [40] as a part of sva package [41], *Quantile Normalization (QN)* [42], *Remove Unwanted Variation (RUV)* [43] and *Harman* [44]. Currently, the most common technique to remove the systematic batch effects from biological data is *ComBat* [40] which based on empirical Bayes method to estimate batch effects and to adjust data across genes. However, often a major limitation of its use is missed: the studies merged have to be more or less balanced with respect to the case-control breakdown of samples. If it is not the case, some biological heterogeneity can vanish [36]. The breakdown by phenotype for each study we use in our work is represented in the Table 1 and shows presence of some imbalance in outcomes. Therefore, we wanted to additionally examine some other normalization methods comprehensively.

The *Quantile Normalization* technique is the same method used for normalization at the probe level, but as it was shown before [42] it can be applied for batch effect elimination as well.

The *RUV* method [43] is based on adjustment on so-called negative control genes, that are expected not to be differentially expressed across phenotypes. There is still an open question which set of genes would be most suitable for this role. Basically, so far there are two possible ways to obtain a list of control genes: housekeeping genes and empirically found ones. The housekeeping genes are defined to be expressed on the similar level across all tissues and involved in basic cell processes [104]. However, besides the fact that there is no fully established list of housekeeping genes [105], it was shown that they might not be the best to represent negative controls in adjustments [43] since they can be differentially expressed in some tissues [33] or related to diseases [106]. Another method is to empirically find genes that are expressed steadily across merging studies. However, there is some freedom in setting a number of such genes. Too small number can be not enough for proper adjustment, too large – can include some genes important for a current study. Some dependence on studies involved is presented in the search for negative control genes. For some discussion of advantages and disadvantages of both methods see reference [107]. Another challenging factor in applying the RUV method is the parameters adjustment. Depending on implementation of the algorithm, there are two main parameters to adjust: the dimension of the unwanted systematic noise and the ridge smoothing parameter. To find an optimal set of parameters is actually a tricky problem. In our study, we examined three different implementations of RUV method in R packages RUVcorr [107], RUVnormalize [108], and RUVSeq [109] (RUVg function). The first two packages implement naïve RUV-random method which is a variation of the RUV-2 method originally described in [43]. The third method RUVg within RUVseq package is originally designed as a discrete version of RUV-2 methods for RNA-seq counts data. We used this method for performance comparison with other approaches. We compared the performance of these RUV implementations with the housekeeping and empirical negative control genes. We also varied the parameter of the noise dimension *k* but set the smoothing ridge parameter to 0.001 for RUVnormalize and 0 for RUVcorr.

Another promising normalization method we examined is *Harman* [44]. This method is a Principal Component Analysis based optimization technique that maximizes batch removal but keeps some probability of the overcorrection as a parameter. We compared the method performance with the overcorrection parameter set to 0.95.

We considered three types of benchmarks to perform the comparison for normalization methods. We computed the percentage of variability in first ten principal components that can be explained by batch (i.e. by dataset study) and kept the maximum value among those ten as one of three benchmarks. For the other two we used the R package gPCA [110] to compute guided principal components (i.e. the principal components of batch modified data) and obtained a p-value (i.e. the probability of having batch effect in data) and delta statistic (i.e. the ratio of guided and unguided first principal component; the lower the better). Since all three metrics are in the range from 0 to 1, we summed them up and normalized to one and used as a final metric to justify the normalization method performance. The results are represented in the Supplementary Fig S6.

We found the Harman normalization to outperform all methods followed by ComBat and RUV implementations in RUVcorr and RUVseq with housekeeping genes at the noise dimension parameter *k* = 15-17. Further comparison analysis showed that while the data seems batch free, RUV and Harman adjusted a bit too much leaving minimum of biological variability, resulting in much fewer differentially expressed genes in comparison to ComBat. Therefore, we decided to use ComBat for cross-study normalization to keep as much heterogeneity as possible while successfully adjusting for batch effects. We observed that the results of the normalization performance vary depending on datasets: their number, platform, processing methods. There is no one-fit-all technique that should be used blindly to correct for batch effects when merging data. Therefore, it should be advised to perform such comparisons of normalization methods each time performing meta-analysis to choose best among available methods [35].

